# Understanding the impact of crosslinked PCL/PEG/GelMA electrospun nanofibers on bactericidal activity

**DOI:** 10.1101/322321

**Authors:** Mirian Michelle Machado De-Paula, Paria Ghannadian, Samson Afewerki, Fernanda Roberta Marciano, Bartolomeu Cruz Viana, Samarah Vargas Harb, Nicole Joy Bassous, Thomas Jay Webster, Anderson Oliveira Lobo

## Abstract

Herein, we report the design of electrospun ultrathin fibers based on polycaprolactone (PCL), polyethylene glycol (PEG), and gelatin methacryloyl (GelMA), and their potential bactericidal activity against three different bacteria *Staphylococcus aureus, Pseudomonas aeruginosa*, and *Methicillin-resistant Staphylococcus aureus* (MRSA). We evaluated the morphology, chemical structure and wettability before and after UV photocrosslinking of the produced scaffolds. Results showed that the developed scaffolds presented hydrophilic properties after PEG and GelMA incorporation. Our developed scaffolds were thus able to significantly reduce gram-positive, negative, and MRSA bacteria. Furthermore, we performed a series of study for better mechanistic understanding of the scaffolds bactericidal activity through protein adsorption study and analysis of the reactive oxygen species (ROS) levels. In summary, we have demonstrated the design and generation of electrospun fibers with improved hydrophilicity and efficient bactericidal activity without the association of any antibiotics.

## 1. Introduction

Microbial infections are immense health problems, in particular with respect to wound healing.^1^ Here, a microbial infection at the wound site can severely prolong the healing process, leading to necrosis, sepsis, and even death.^2^ The most common pathogenic bacteria found in the hospital environment are *Escherichia coli (E. coli), Pseudomonas aeruginosa (P. aeruginosa), Staphylococcus aureus* (*S. aureus*), *Staphylococcus epidermis (S. epidermis)*, and various filamentous fungi as well as yeasts (*i.e., Candida spp*.).^3-5^ The use of antibiotics can results in an escalating drug resistance in pathogenic microorganisms, which is associated with increased morbidity and mortality and therefore should be avoided when possible.^6^ In fact, according to the World Health Organization (WHO), antibiotic resistance is one of the biggest threats to global health. Therefore, the design of materials with the ability to significantly reduce the administration of antibiotics and avoid infections from highly antibiotic-resistant bacteria in the hospital environment are highly desirable. In this context, degradable polymers with tunable mechanical, biological and chemical properties and ease of fabrication can be attractive for aforementioned applications.^7^ Through the combination of several biomaterials with inimitable properties, a tailor-made hybrid material can be designed without the use of any complicated fabrication approaches or chemistry. However, one requirement is that such materials are compatible with each other avoiding interference or inhibition due to mismatched chemistry or structure, but instead promote properties synergistically.

Coupled with the above, biomaterial nanotexturing has recently been shown to reduce microorganism attachment and growth. One method for the preparation of ultrathin biomaterial fibers can be accomplished by employing electrospinning technology.^8-12^ This approach is simple, efficient, reproducible, cost-efficient, and provides high yield.^13-16^ Here, we envision by merging three polymeric biomaterials (polycaprolactone (PCL), polyethylene glycol (PEG) and photocrosslinkable gelatin methacryloyl (GelMA)) for the generation of ultrathin electrospun fibers, bactericidal property can be obtained. These materials were selected based on their unique properties. Polycaprolactone (PCL) is one of the most extensively employed polyesters in biomedical and tissue engineering applications due to its biocompatibility, mechanical strength, viscoelasticity and biodegradability.^17, 18^ Nevertheless, its hydrophobic property and long degradation time limits its applications. One way to solve such limitations is by merging PCL with other biomaterials to improve its properties such as hydrophilicity, degradability, and controllable mechanical properties.^19-22^ Due date, several strategies have been applied to combine PCL with other polymers to obtain partial or total hydrophilicity to open numerous possibilities for biomedical purposes.^23-30^ Furthermore, PEG is biocompatible, hydrophilic and has the ability to reduce protein adsorption.^31^ However, it still has limits in that it cannot be used alone to produce mats, but rather needs to be combined with other polymers.^32, 33^

Moreover, GelMA is a biomaterial that resembles the natural components in the extracellular matrix in native tissues and displays several biological significance properties such as biocompatibility, biodegradability, low antigenicity and promoting cell adhesion due to the RGD (Arg-Gly-Asp) motif. Nevertheless, its poor mechanical properties and fast enzymatic degradation have limited its biomedical applications and in this context, lacks bactericidal properties.^34^,^35^ Previously gelatin has been reported to be a good source for bacteria growth.^36^

Herein, we electrospun a combination of PCL, PEG and GelMA and evaluated the potential of the resulting UV crosslinked scaffolds as bactericidal biomaterial and the understanding about a possible mechanism of reduction using specific biological assays. To the best of our knowledge, this is the first report on the electrospinning of the combination of PCL, PEG, and GelMA, evaluating their bactericidal activity against reduce *S. aureus, P. aeruginosa,* and *MRSA* growth. In doing so, this study has opened numerous perspectives to use a new bactericidal scaffolding system for a wide range of biomedical and tissue engineering applications.

## 2. Materials and methods

### Materials

The following chemicals were purchased from Sigma-Aldrich (St. Louis, MO, USA): PCL (molecular weight (Mw) 80,000), PEG (Mw 8,000), Gelatin (Type A, 300 bloom from porcine skin), methacrylic anhydride (MA), dimethyl sulfoxide (DMSO), and photoinitiator 2- hydroxy-1-[4-(hydroxyethoxy) phenyl]-2-methyl-1-propanone(Irgacure 2959). Hexafluoroisopropan-2-ol (HFIP) was purchased from Oakhood Chemical. Dulbecco’s phosphate buffered saline (DPBS), trypsin-EDTA (ethylenediaminetetraacetic acid) and penicillin-streptomycin were purchased from Gibco (MD, USA). Alpha-modified Eagle’s medium (Alpha-MEM) was supplied by Invitrogen (Grand Island, NY, USA). HyClone characterized fetal bovine serum (FBS) and pre-cleaned microscope slides were obtained from Fisher Scientific (Waltham, MA, USA). *S. aureus, P. aeruginosa*, and *Methicillin-resistant S. aureus (MRSA)* were acquired from the ATCC strain 12600. The UV source (Omnicure model S2000) was provided by EXPO Photonic Solutions Inc. (Ontario, Canada).

### Procedure for the synthesis of GelMA

GelMA was prepared following our previously reported protocol (**Fig. 1a**).^37^ Shortly, gelatin type A (10 g) in 100 mL PBS was stirred at 50 °C until it was fully dissolved. Afterwards, methacrylate anhydride (3 mL) was added and stirred for additional 2 h. The material was purified through dialysis tubing (12–14 kDa MWCO, spectrum Lab Inc.) for 7 days at 40 °C. Finally after freeze-drying GelMA was obtained as a white solid.

**Fig. 1.**
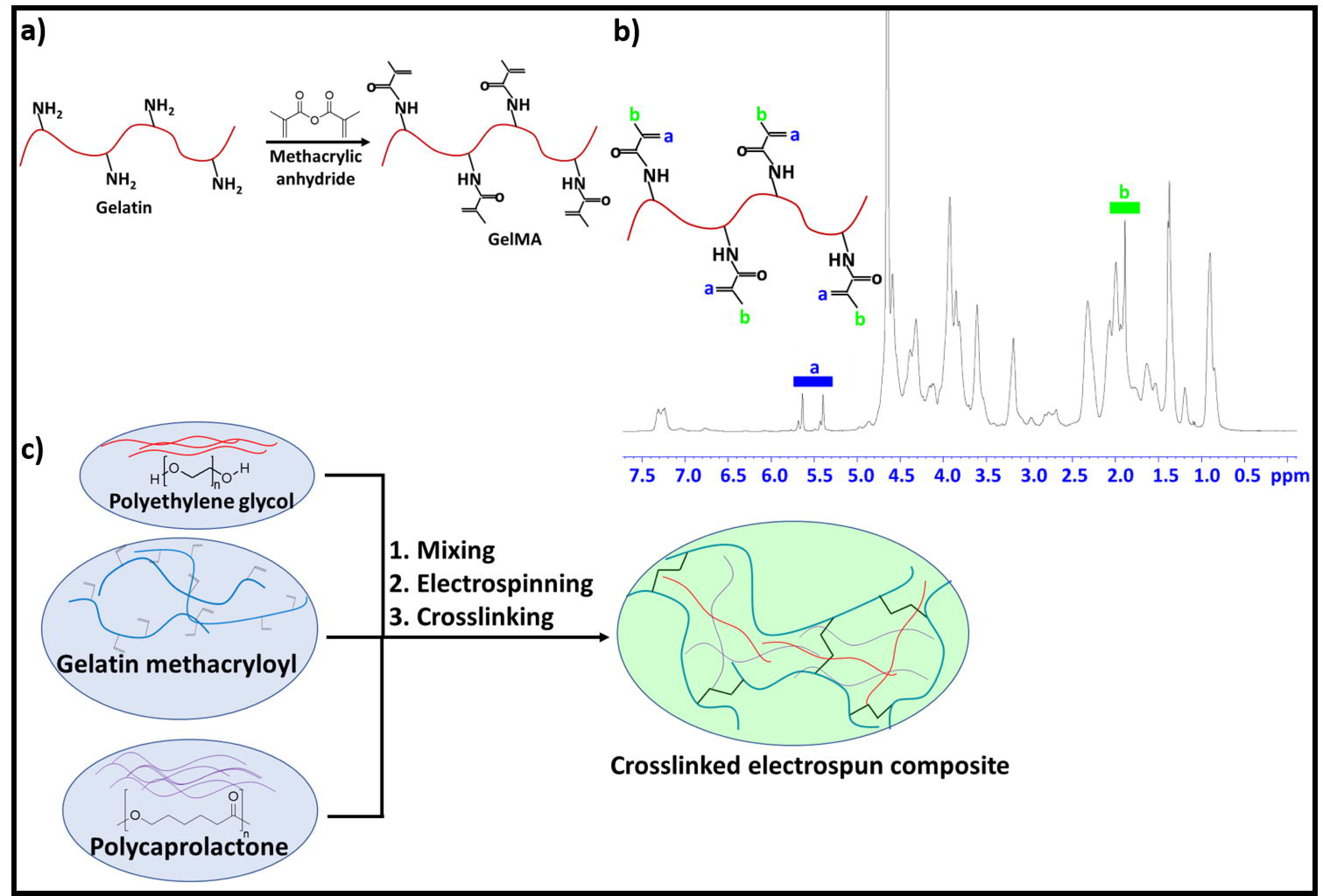
a) The synthetic route for the preparation of gelatin methacryloyl (GelMA). b) ^1^H-NMR of the prepared GelMA identifying the specific groups. c) The fabrication approaches for the preparation of the PCL, PEG and GelMA combined electrospun fiber.

### ^1^H NMR analysis

Prior NMR running, the GelMA samples were dissolved in deuterium oxide (D_2_O) at a concentration of 30 mg/mL and the running was performed at 37 °C. The data were processed using ACD LABS 12.0 software. The degree of substitution of GelMA was determined to 71% by NMR analysis (**Fig. 1b**).

### Preparation of the solutions prior electrospinning

The samples were prepared separately by mixing PCL (0.24 g), PEG (0.15 g) and GelMA (0.05 g) in 5 mL of HFIP for each sample in vials. Subsequently, the vials were closed and dissolved under stirring at room temperature for 24 h. Then, the solutions were placed in a syringe (BD Yale, 3 mL) followed by connecting a needle to the syringe (Inbras, 23G). The electrospinning was carried out at 17 kV (Nanospinner Machine, Inovenso) as a positive voltage. An aluminum foil collector plate (20 cm × 20 cm square plate) was used as the anode, a needle-collector distance of 10 cm was used, and 1 mL/h was used as the solution flow rate (Harvard, PHD 2000). The temperature and humidity were controlled at 21–23 °C and 44–54%, respectively.

### Photocrosslinking of the produced scaffolds

The photocrosslinking solution was prepared by adding 1.0 g of photoinitiator to 10 mL of ethanol in the absence of the light and stirred until completely dissolved. The uncrosslinked electrospun mats was first immersed in a glutaraldehyde (25% in deionized (DI) water) solution (10 mL/500 mL ethanol) overnight and then extensively washed with a solution of glycine (3.75 g in 250 mL of DI water). Subsequently, the mats were immersed in an ethanol solution containing photoinitiator (10 wt%) for 2 h and then exposed to a 365-nm UV light (OminiCure^®^- 2000 Series) at a 10-cm working distance for 10 min. Then, the crosslinked mats were washed 3 times with ethanol. Afterwards, the mats were washed in DI water and dried under vacuum. **Fig. 1c** illustrates a possible mechanism of PCL:PEG:GelMA interaction after photocrosslinking.

### Characterization of the produced scaffolds

A scanning electronic microscope (FEI Quanta 200, 3 kV) was used to analyze the scaffold morphology. A thin gold layer was evaporated onto all scaffold surfaces before analysis to improve image acquisition.

Fourier transform infrared spectrometry (FTIR, Spotlight-400, Perkin Elmer FTIR Imaging System) was used to analyze the chemical groups in the biomaterial before and after photocrosslinking under a controlled atmosphere and transmittance mode. The spectra were collected at regions between 4000–400 cm^−1^ and were analyzed in three different regions. The regions of interest were analyzed, and their vibrational modes indexed.

For wettability analysis, the contact angle (CA) between the surface of the scaffold and a DI water or diiodomethane drop was monitored in the dynamic mode, using a contact angle device (Krüss, Model DSA 100). Before analysis, a careful calibration was performed in accordance to the equipment manual and all the scaffolds were fixed on a Teflon base. Then, DI water and diiodomethane (2 μL) were dropped onto the scaffold surfaces and measured after 1 s. The atmosphere and humidity (∼ 60%) were controlled during all the measurements. The surface tension and adhesion force were calculated and described in the supplementary data.

### Bacteria tests

*S. aureus* (ATCC 25923), *P. aeruginosa* (ATCC 25668), and *Methicillin-resistant S. aureus (MRSA)* (ATCC 43300) were inoculated in 3% Tryptic soy broth (TSB) and after 14 h, the bacteria were plated at a concentration of 10^4^ cells/mL onto each sample (1 cm × 1 cm). After 24 h, the samples were lightly rinsed with PBS (x2) to remove any non-adherent bacteria. This bacterial solution was then diluted serially (10X; 100X; 1000X) and plated on tryptic soy agar plates and incubated overnight (∼14 h). Afterwards, the colonies were counted to calculate the colony-forming units (CFUs). The data were normalized by raw PCL data to highlight the bactericidal activity of the produced scaffolds without the use of antibiotics.

### Protein adsorption

For protein adhesion test, the fibers were placed in a 24-well plate, 0.5 mL of TSB or casein solution were added and the plate was kept inside an incubator at 37 °C for 24 h. After this time, the samples were transferred to a new plate, washed twice with PBS and 0.5 mL of immunoprecipitation assay (RIPA) buffer was added to each well. After 10 min in contact with RIPA buffer, the solution inside the wells was mixed with pipette to ensure homogeneity. Using a 96-well plate, 25 μl of the samples were mixed with 200 μl of working reagent from Pierce^TM^ bicinchoninic acid (BCA) Protein Assay Kit, Thermo Scientific. The plate was placed on a plate shaker for 30 s and then incubated at 37 °C for 30 min. After incubation, the plate was coolED to room temperature and the absorbance was measured at 562 nm using a spectrophotometer (SpectraMax® M3, Molecular Devices).

### Analyses of the oxygen species

The reactive oxygen species (ROS) levels before and after UV crosslinking was analyzed using Fluorimetric Hydrogen Peroxide (H_2_O_2_) Assay Kit (Sigma-Aldrich) and the Cellular ROS/Superoxide (O_2_^−1^) Detection Assay (Abcam). 1 mL of *S. aureus, P. aeruginosa* and *Methicillin-resistant S. aureus (MRSA*) solutions with concentration of 10^4^ CFU/mL were placed into 24-well plates containing the fibers. The plates were kept inside the incubator at 37 °C for 24 h. After this time, the bacteria solution was removed, subsequently, the scaffolds were washed 2x with 1 mL PBS each, and then placed inside an eppendorf tube containing 1 mL of PBS. After vortex for 15 min, the solutions were used to perform the 2 tests.

For the hydrogen peroxide assay, 50 μl of each solution was placed in a 96-well plate and mixed with 50 μl of working solution from the assay kit. The 96-well plate was incubated at room temperature for 30 min followed by measuring the absorbance at 540 nm using a spectrophotometer (SpectraMax^®^.Paradigm^®^, Molecular Devices).

In the more generalized ROS and superoxide detection process, a microplate assay procedure was implemented in which a mixture containing reactive cell permeable probes was added to the test samples. Specifically, 100 μL of each test and control was transferred from the eppendorf tubes after vortexing to an opaque 96-well plate. Positive controls were established in which a set of cells were treated with Pyocyanin in order to induce ROS generation. 100 μL of the ROS/superoxide detection mixture was added to the cell suspensions inside the 96-well plate, and the plate was allowed to incubate at 37 °C for 1 h in the dark. After the staining period, the 96-well plate was loaded onto the spectrophotometer with standard fluorescein (Excitation/Emission = 490/525 nm) and rhodamine (Excitation/Emission = 550/620 nm) filter sets for detecting ROS and superoxides, respectively.

### Statistical analysis

All experiments were conducted in triplicate and were repeated at least three different times with analysis of variance (ANOVA) followed by Student’s t-tests to determine statistical differences between mean values. The populations from the analyzed scaffolds were obtained with normal distribution and independent to each experiment.

## 3. Results

The surface and structural characterization of the scaffolds are summarized in **Fig. 2**. One clearly observation is that ultrathin and microfibrous scaffolds in the absence of any beads were obtained independently of the analyzed groups. When the PCL solution was electrospun, a non-homogeneous behavior, and some defected fibers were produced (**Fig. 2a1**). More details about the surface defects can be seen in **Fig. 2a2**. and **Fig. 2a3**, which highlights the partial roughness structure and defects on the scaffold. The distribution of diameters is depicted in **Fig. 2a4**, where PCL presented microfibrous structures (diameter of 1.53 μm ± 0.74 μm). After the incorporation of PEG, a discrete regularity in the material was obtained (**Fig. 2b1**) and the PCL:PEG scaffolds presented a better smoothness (**Fig. 2b2**). The smooth surface of the fibers could be more easily identified in the magnified **Fig. 2b3** image. The thickness also decreased by 30% compared to PCL (**Fig. 2b4**, 0.92 μm ± 0.43 μm). Furthermore, after the addition of GelMA the electrospun fibers transformed to an even more regular scaffold (**Fig. 2c1**). **Fig. 2c2** illustrates details from the non-defected structures with a homogeneous behavior. By comparing the detailed SEM image **Fig. 2c3**, with PCL (**Fig. 2a1-a3**) and PCL:PEG (**Fig. 2b1-b3**) scaffolds, a smoother surface was obtained. Moreover, the distribution of the diameters from the PCL:PEG:GelMA scaffolds showed ultrathin fibers (0.24 μm ± 0.10 μm) smaller than PCL. The crosslinked PCL:PEG:GelMA scaffolds also presented a more homogeneous surface without any further defects (**Fig. 2d1-d3**), and with increases in diameter (**Fig. 2d4**) after crosslinking compared to non-crosslinked PCL:PEG:GelMA (**Fig. 2c1-c3**).

**Fig. 2.**
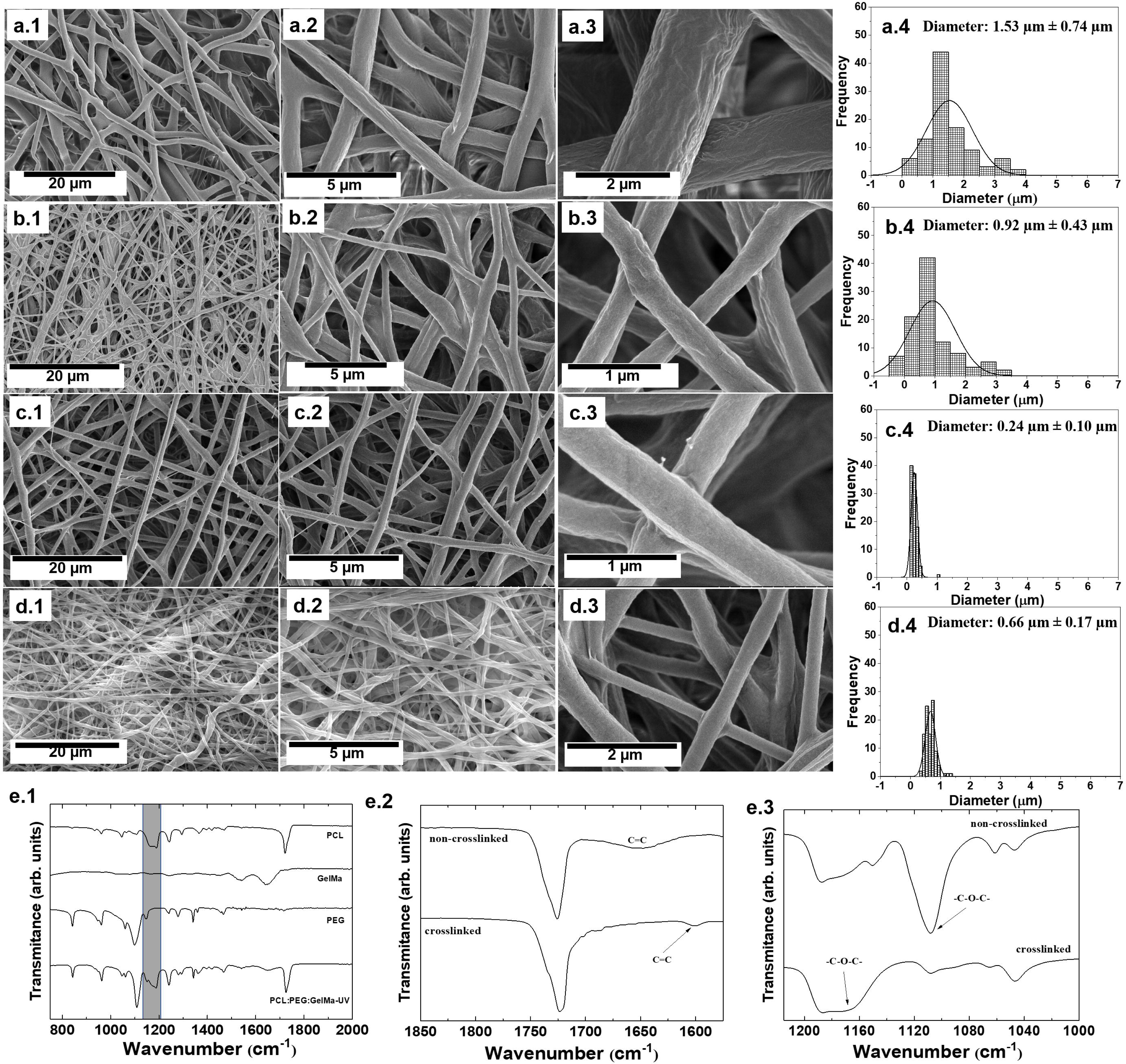
Characterization of the produced scaffolds. Scanning electron microscopy of: (**a1**–**a3**) PCL, (**b1**–**b3**) PCL:PEG, (**c1**–**c3**) PCL:PEG:GelMA and (**d1**–**d3**) crosslinked PCL:PEG:GelMA scaffolds. Distribution of diameters from (**a4**) PCL, (**b4**) PCL:PEG, (**c4**) PCL:PEG:GelMA and (**d4**) crosslinked PCL:PEG:GelMA scaffolds. (**e1**–**3**) Corresponding FTIR spectra of the produced scaffolds. (**e1**) Details of all electrospun scaffolds identifying the main peaks and bands to each polymer. (**e2**) The specific FTIR region identifying the vibrational region referred to the C=C bond from non- and crosslinked scaffolds. (**e3**) The main FTIR region containing the vibration of ether groups related to the methacryloyl substitution of GelMA. The groups were analyzed before and after UV irradiation.

The structural analysis and chemical interactions were recorded on the FTIR analysis (**Fig. 2e1-e3**). The main peaks for each separate polymer are shown in **Fig. 2e1**. In the FTIR from the PCL, the main peaks could be identified at: 731 cm^−1^ (-(CH_2_)_*n*_), 1157 cm^−1^(C-O and C-C stretching in the amorphous phase), 1170 cm^−1^ (symmetric C-O-C stretching), 1190 cm^−1^ ((*v*(O-C-O)), 1240 cm^−1^ (asymmetric C-O-C stretching), and 1342 cm^−1^ (CH_2_) ^38, 39^. Also, the fingerprint region center was observed at: 1637 cm^−1^ (amide I), 1529 cm^−1^ (amide II), and 1448 cm^−1^ (amide III), corresponding to the stretching of the C=O bond, bending of the N-H bond, and plane vibration of C-N and N-H, respectively ^40^. Meanwhile, the main peaks related to PEG are: 839 cm^−1^ (δ(C–H)), 951.26 cm^−1^ (CH_2_), 1115.19 cm^−1^(C-O), and 1350 cm^−1^ (*v* (C — C)) ^41, 42^. Inthe region 1100–1200 cm^−1^ of the mixed polymers, the intermolecular forces such as hydrogen bonds, ions, and charge transfers could be observed ^43^.

**Fig. 2e2** and **2e3** show the difference between non- and crosslinked scaffolds. It is known that FTIR helps to analyze the fraction of unreacted C=C double bonds of methacrylate or methacrylamide groups ^44^. The band centered at 1634 cm^−1^ indicating the presence of the double bond was absent after UV irradiation (**Fig. 2e2**), whilst this was present on the scaffold before UV irradiation (identified at **Fig. 2e2**). The increased chemical crosslinking resulted in an enhanced physical network ^44, 45^.After UV irradiation, a peak centered at 1600 cm^−1^ appeared (acrylic double bonds, C=C) and at the same time, the peak at 1100 and 1170 cm^−1^ disappeared (**Fig. 2e3**, related to the -C-O-C- vibration commonly found when scaffolds contains GelMA). This can be associated to the reaction of the methacryloyl moiety during crosslinking ^43, 46^.

The bactericidal effect was evaluated against three most prominent bacteria in the hospital environment, such as: *S. aureus* (**Fig. 3a**), *P. aeruginosa* (**Fig. 3b**), and *MRSA* (**Fig. 3c**). We compared these results to raw PCL. After UV crosslinking, the scaffolds inhibited the growth of *S. aureus* (**Fig. 3a**, gram-positive) compared to raw PCL. Interestingly, the PCL:PEG:GelMA-UV scaffolds reduced *P. aeruginosa* by 10-fold comparable to the material without UV crosslinking (**Fig. 3b**). Moreover, our developed scaffolds showed high antibacterial activity against multiple drug-resistant *S. aureus* bacteria (**Fig. 3c**, MRSA) and the same efficiency was observed for the materials before and after UV crosslinking. We measured the contact angle (CA) from our developed scaffolds to explain this bactericidal activity presented for the PCL:PEG:GelMA before and after UV crosslinking and calculated their surface energy and adhesion force of bacteria attached using thermodynamic approaches (see supplementary information). Interesting, the PCL:PEG:GelMA-UV presented a CA (water, 69°) higher than the same material without UV crosslinking (water, 25°), probably due to a more dense material obtained after crosslinking introducing covalent bonds to the material. However, the CA using diiodomethane gave the same result for both of the materials with/without crosslinking (∼5°, supplementary information, **Table S2**). Then, we calculated the surface energy of our developed scaffolds and the material after UV crosslinking promoting the bactericidal activity showed a lower surface energy (53.3 mJ/m^2^) compared to the material without UV crosslinking (71.4 mJ/m^2^). Moreover, the adhesion force was calculated using a thermodynamic approach where the average value of work obtained for the adhesion of bacteria from PCL:PEG:GelMA were Δ*F*_*Adh*_= −34.6 mJ/m^2^ (without crosslinking) and Δ*F*_*Adh*_ = −22.6 mJ/m^2^ (after crosslinking). These results indicated that the scaffold surfaces before UV crosslinking was more favorable for bacteria adhesion than after ^47^.

**Fig. 3.**
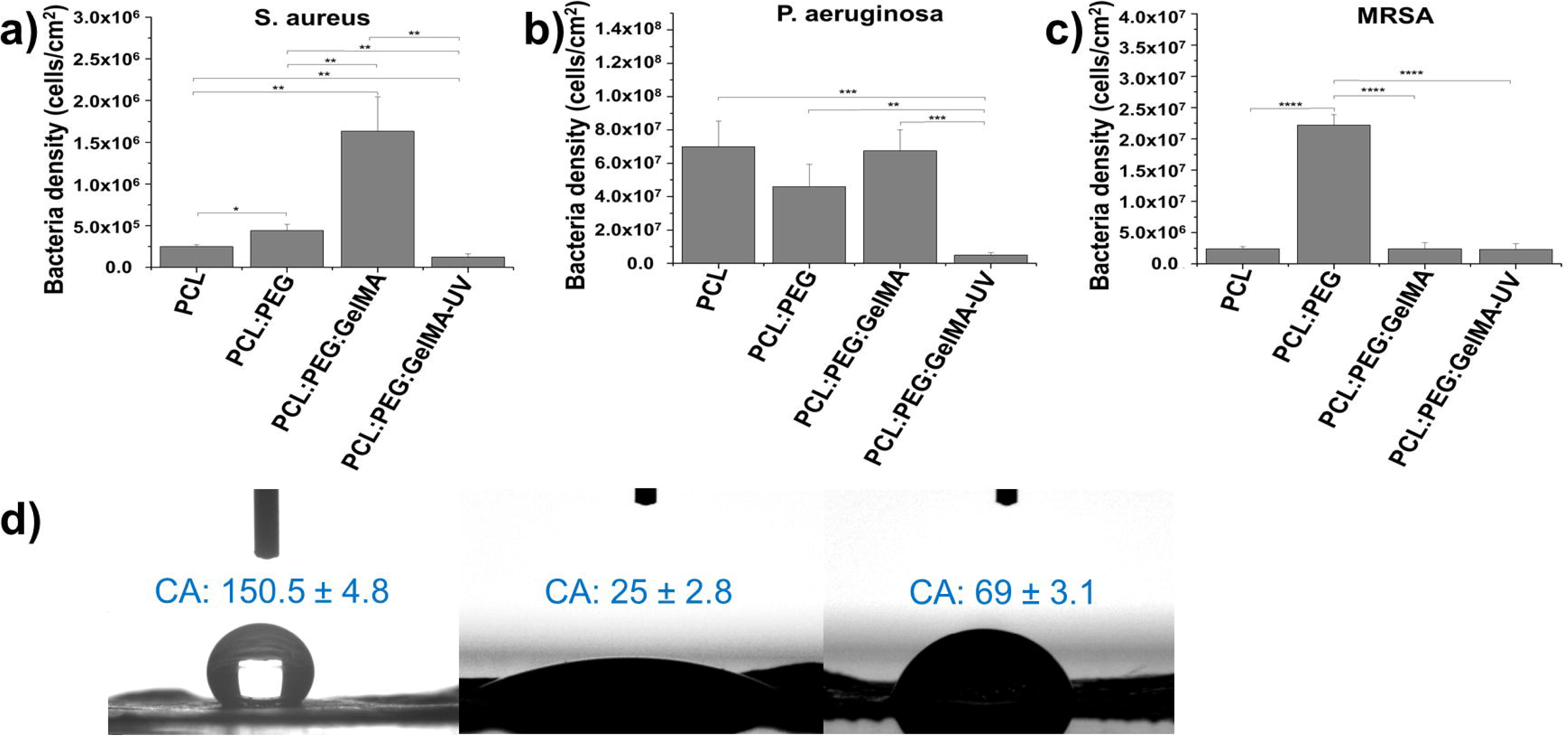
Bactericidal study on the electrospun scaffolds after 24 h (**a**) *S. aureus* growth reduction.(**b**) *P. aeruginosa* growth reduction. (**c**) MRSA growth reduction. The data were plotted in percentage and compared to raw PCL. N=5. (**) p< 0.01, (***) p<0.001 and (****)p< 0.0001 mean statistical differences. (**d**) The wettability measured through contact angle analysis from water on PCL, PCL-PEG-GelMA and PCL-PEG-GelMA-UV scaffolds.

In order to analyze the effect of the UV crosslinked fiberson the bactericidal properties, the non-crosslinked and crosslinked fibers were submitted to protein adsorption assay (**Fig. 4a),** hydrogen peroxide assay (**Fig. 4b)**, reactive oxygen species (ROS) assay (**Fig. 4c)** and superoxide assay (**Fig. 4d)**. The protein adsorption assay showed for both TSB and casein an increase of the protein concentration adsorbed onto the crosslinked fibers. Impressively, bacterial adhesion to the implant surfaces is inhibited, despite the generalized understanding that TSB and casein aid in bacterial growth ^[34]^. The reason for this is that casein adsorption to an implant surface prevents biomolecular attachment, and the increased protein adsorption over a restricted time period correlates with antibacterial properties due to short-scale bacterial adhesion dynamics ^[35]^. From this study, it is possible to conclude that the UV crosslinking of scaffold composites improves protein adsorption, and, correspondingly, reduces bacterial attachment and growth.

**Fig. 4.**
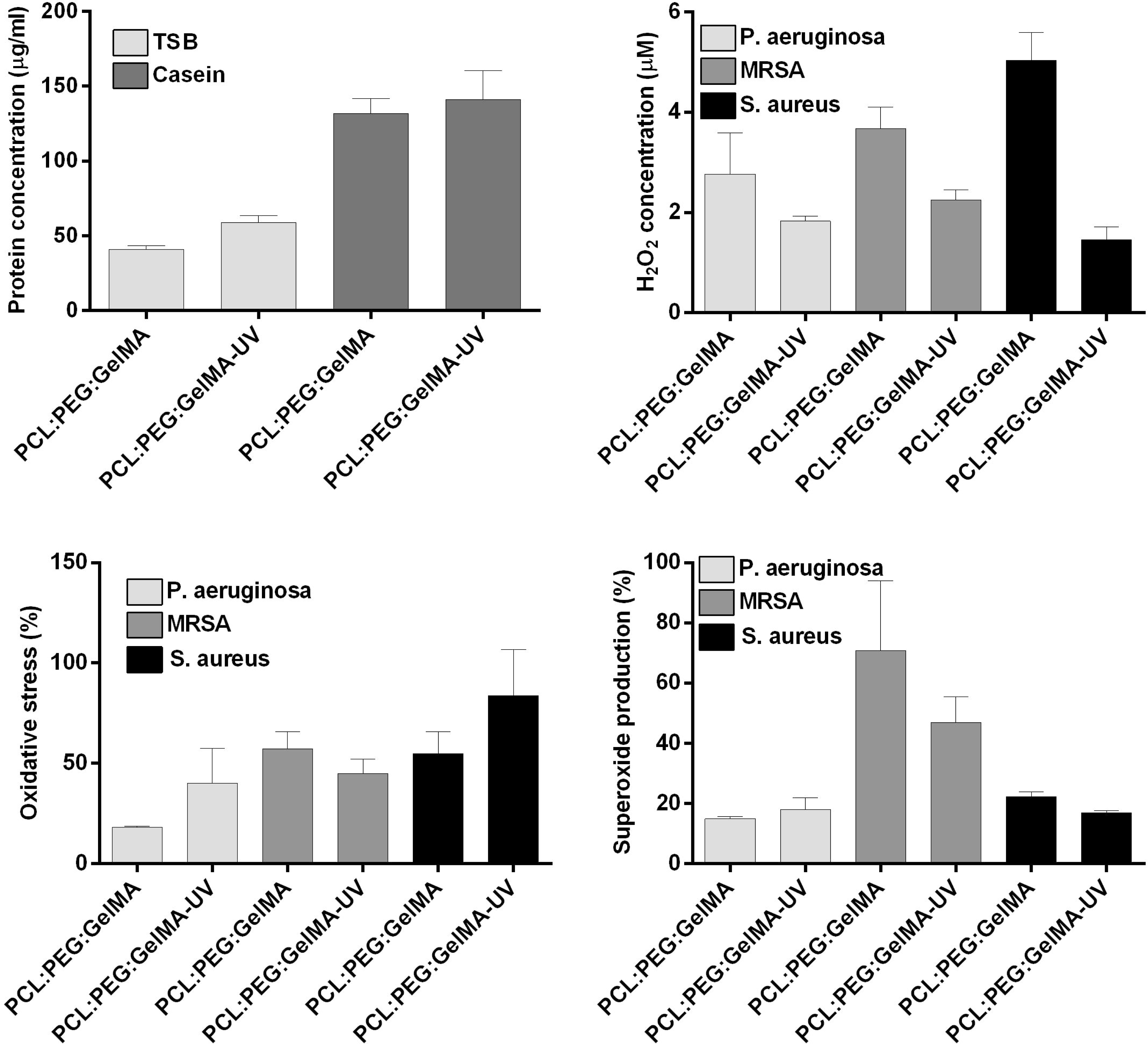
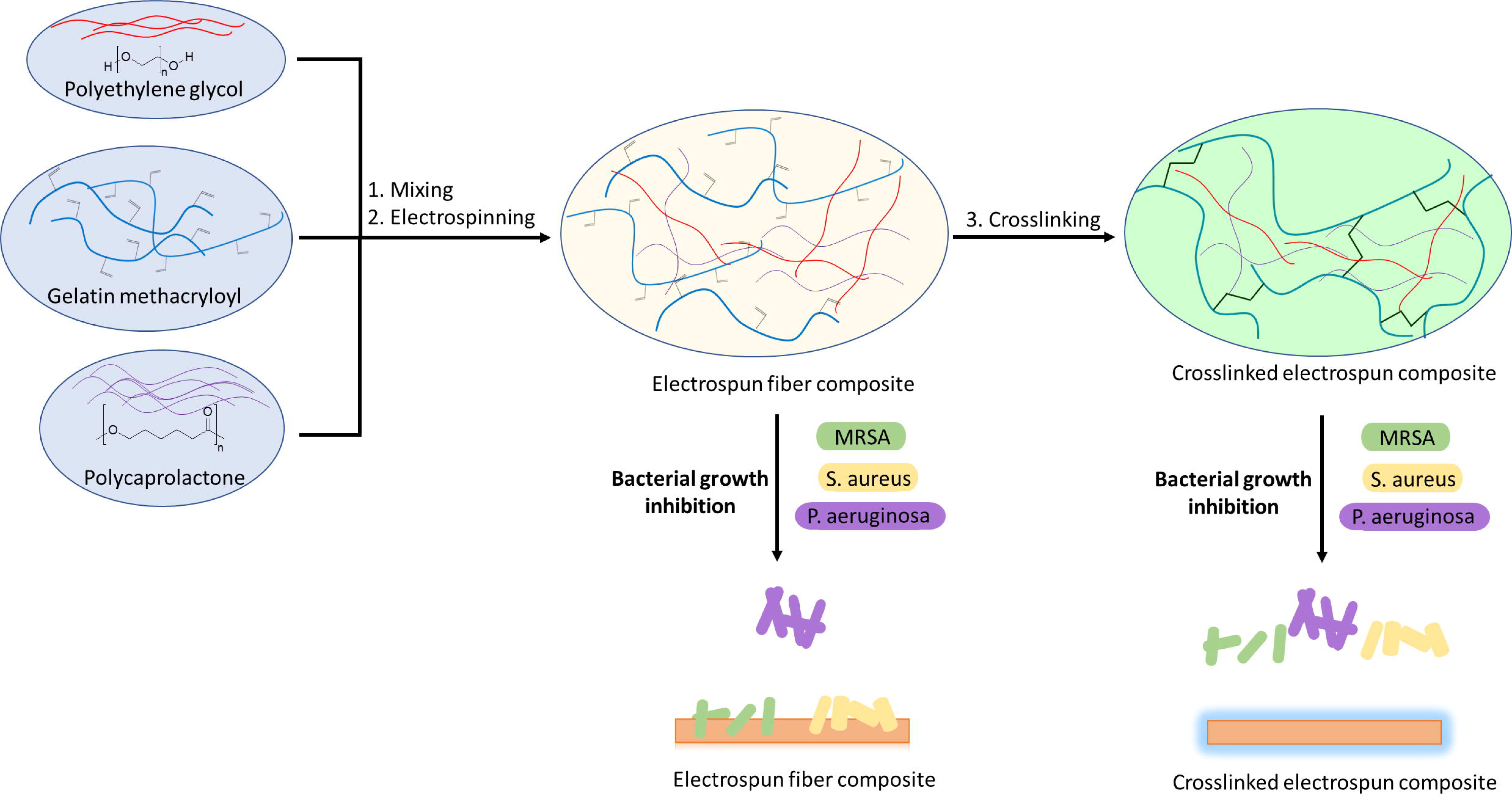
(**a**) Protein adsorption after 24 h. (**b**) Hydrogen peroxide assay, (**c**) reactive oxygen species (ROS) assay and (**d**) superoxide assay, after 24 h immersed in *S. aureus, P. aeruginosa* and MRSA bacteria solution, where percentages are given relative to positive control cells treated using a Pyocyanin ROS inducer. N=3.

The UV crosslinking also influences the production of reactive oxygen species, such as hydrogen peroxide (**Fig. 4b**). The data obtained using the Fluorimetric Hydrogen Peroxide Assay Kit indicates that crosslinking of the fibers contributed to a decrease in the production of H_2_O_2_ for all three of the bacterial types evaluated. Nevertheless, the overall oxidative stress imposed on the *P. aeruginosa* and *S. aureus* strains due to the combined effect of H_2_O_2_, hydroxyl radicals (HO·), peroxynitrite (ONOO-), and peroxy radicals (ROO·), as measured using the ROS/Superoxide Detection Assay, is augmented for the UV crosslinked samples (**Fig. 4c**). This indicates that scaffold crosslinking plays a role in promoting the production of ROS that facilitate the self-destruction of *P. aeruginosa* and *S. aureus* cells. This conclusion, however, cannot be drawn for the drug-resistant MRSA strain. Indeed, a modest reduction in the ROS levels for MRSA suspensions that were grown along UV-crosslinked scaffolds was documented. Moreover, an increase in the overall oxidative stress imposed on the *P. aeruginosa* and *S. aureus* strains due to scaffold crosslinking did not correlate with an increase in superoxide production (**Fig. 4d**), and the amount of (O_2_^−1^) generated by these bacteria upon exposure to both the UV-crosslinked and non-crosslinked samples was approximately the same. A significant decrease inthe production of O_2_ by MRSA cells incubated on the crosslinked scaffolds, relative to the non-crosslinked scaffolds, however, was observed.

## 4. Discussion

Herein, we combined three different polymers to obtain ultrathin electrospun meshes. To date, we are not aware of any prior reports that have successfully demonstrated the electrospinning of the combination of PCL, PEG, and GelMA and evaluation on their bactericidal activity. Their efficiency was more evident after UV photocrosslinking, which could be associated to the closer interaction between the polar groups (such as -NH_2_, -OH, and the carbonyl group (-CO)) in GelMA and -OH found on the PEG chain, and also due to the more compact scaffold obtained. The surface energy change of the scaffolds after UV crosslinking (**Table S2**) can lead to changes in protein adsorption and bioactivity altering cell functions and, at the same time, reducing bacteria adhesion and proliferation.^48^ Furthermore, it is widely accepted that reactive chemical groups on micro and nano-fibers can disrupt bacteria membranes via oxygen free radical formation. Therefore, it can be inferred, especially in the case of MRSA, that factors besides ROS and superoxide production contribute to the bactericidal efficacy of PCL:PEG:GelMA scaffolds that are crosslinked. This is beneficial considering that cellular mechanisms have evolved to evade killing by self-generated ROS (**Fig. 4c**). For instance, adhesion prevention appears to factor significantly into disabling MRSA attachment and growth along scaffolds that incorporate GelMA.

Scaffolds based on different amounts of PCL, GelMA, and gelatin have been reported by Correa et al. ^49^ However, the authors first produced PCL mats and then incorporated GelMA and gelatin afterwards using *in situ* coating. The process presented here is different and more interesting to tissue engineering applications since we electrospun PCL, PEG and GelMA together and the subsequent photocrosslinked composite promoted bactericidal activity. A more hydrophilic scaffold (CA ∼25°) was obtained after GelMA and PEG inclusion. The potential of water sorption of GelMA and GelMA:PLGA mats were also investigated by Zhao *et al*. ^50^. The authors obtained a highly porous scaffold which increased the diameter to ∼70% after 24 h.

Other authors have explored the influence of GelMA and PEG to induce vascularization and extracellular matrix calcification ^51-57^, however, the bactericidal properties have not yet been investigated.

Anabi *et al*. reported that hydrogels based on GelMA conjugated with an amphiphilic peptide reduced bacteria growth ^58^. However, the bactericidal effect against *S. aureus* was only observed after peptide incorporation. Commonly, micro- and nanoparticles have been associated to short sequences of cationic amino acids to possess a broad-spectrum of bactericidal activity against Gram (+/-) bacteria ^59-62^. Here, impressively, *S. aureus* and *P. aeruginosa* were reduced to a half amount compared to raw PCL without requiring additional steps of peptide functionalization or antibiotic incorporation. Additionally, no statistical difference was observed compared to PCL:PEG:GelMA before and after UV against MRSA.

Rujitanaroj *et al*. combined gelatin and GelMA with silver nanoparticles and evaluated their bactericidal effect ^63^; the use of silver has recently been questioned due to high mammalian cell toxicity. Moreover, the authors only observed the bactericidal effect when silver nanoparticles were added. In this manner, it is plausible to at least preliminarily compare the antimicrobial mechanism of our scaffold to those obtained when amphiphilic peptides (AMPs) were used. It is noteworthy that our developed scaffold had no positive charge in its structure, indicating that the binding interactions between the hydrogel and gram-positive and negative bacteria were unlikely to be caused by electrostatic interactions, as seen in AMPs, which involved bacterial cell wall and membrane disruption.^64-66^ We believe that the design of a more hydrophilic scaffold promoted the promising results obtained from the bacteria study by altering initial protein adsorption that bacteria depend upon to attach ^67^. All these characteristics highlight the application of PCL:PEG:GelMA as novel scaffolds for tissue engineering purposes. Further studies will be needed to determine the exact mechanisms (including initial ion/protein adsorption) responsible for the reduced bacteria growth on the present scaffolds.

## 5. Conclusions

In this report we disclose, for the first time, the preparation and bactericidal proficiency investigation of electrospun scaffolds by merging PCL:PEG:GelMA polymers. The designed ultrathin fibers scaffolds were hydrophilic comparable to raw PCL. Moreover, the developed scaffolds were also able to efficiently prevent the growth of *S. aureus, P. aeruginosa*, and MRSA after UV crosslinking. In conclusion, all these findings suggest that the designed biomaterial is a promising candidate for various biomedical applications.

## Acknowledgements

The authors would like to thank the Coordination for the Improvement of Higher Education Personnel (CAPES, grant numbers AOL#88881.120138/2016-01 and FRM#88881.120221/2016-01), Brazilian National Council for Scientific and Technological Development (CNPq, AOL#303752/2017-3 and FRM#304133/2017-5) and Universidade Brasil for scholarships. The authors also acknowledge Northeastern University for funding this study. S. A. gratefully acknowledges financial support from the Sweden-America Foundation (The family Mix Entrepreneur foundation), Olle Engkvist byggmästare foundation and Swedish Chemical Society (Bengt Lundqvist Memory Foundation) for a postdoctoral fellowship

